# A mathematical model of flow-mediated coagulation identifies factor V as a modifier of thrombin generation in hemophilia A

**DOI:** 10.1101/495374

**Authors:** Kathryn G. Link, Michael T. Stobb, Matthew G. Sorrells, Maria Bortot, Katherine Ruegg, Marilyn J. Manco-Johnson, Jorge A. Di Paola, Suzanne S. Sindi, Aaron L. Fogelson, Karin Leiderman, Keith B. Neeves

**Affiliations:** Department of Mathematics, University of Utah, Salt Lake City, UT, USA; Department of Applied Mathematics, University of California, Merced, Merced, CA, USA; Department of Chemical and Biological Engineering, Colorado School of Mines, Golden, CO, USA; Department of Bioengineering, University of Colorado, Denver | Anschutz Medical Campus, Aurora, CO, USA; Hemophilia and Thrombosis Center, University of Colorado, Denver | Anschutz Medical Campus, Aurora, CO, USA; Department of Pediatrics, University of Colorado, Denver | Anschutz Medical Campus, Aurora, CO, USA; Department of Biomedical Engineering, University of Utah, Salt Lake City, UT, USA; Department of Applied Mathematics and Statistics, Colorado School of Mines, Golden, CO, USA

**Author notes:** KGL, MTS, MGS and MB contributed equally to this work.

**Keywords:** hemophilia A, hemostasis, factor V, sensitivity analysis

## Abstract

Hemophilia A is a bleeding disorder categorized as severe, mild, and moderate deficiencies in factor VIII (FVIII). Within these categories the variance in bleeding severity is significant and the origins unknown. The number of parameters that could modify bleeding are so numerous that experimental approaches are not feasible for considering all possible combinations. Consequently, we turn to a mathematical model of coagulation under flow to act as a screening tool to identify parameters that are most likely to enhance thrombin generation. We performed global sensitivity analysis on 110,000 simulations that varied coagulation factor levels by 50-150% of their normal values in humans while holding FVIII levels at 1%. These simulations identified low factor V (FV) levels as the strongest candidate, with additional enhancement when combined with high prothrombin levels. This prediction was confirmed in two experimental models: Partial FV inhibition boosted fibrin deposition in flow assays performed at 100 s^−1^ on collagen-tissue factor surfaces using whole blood from individuals with mild and moderate FVIII deficiencies. Low FV (~50%) or partial FV inhibition also augmented thrombin generation in FVIII-inhibited or FVIII-deficient plasma in calibrated automated thrombography. These effects were amplified by high prothrombin levels in both experimental models. Our mathematical model suggests a mechanism in which FV and FVIII compete to bind to factor Xa to initiate thrombin generation in low FV, FVIII-deficient blood. This unexpected result was made possible by a mechanistic mathematical model, providing an example of the potential of such models in making predictions in complex biological networks.

## Introduction

Hemophilia A is a genetic bleeding disorder caused by a deficiency in coagulation factor VIII (FVIII), a protein in blood plasma necessary to generate stable blood clots. FVIII deficiency prevents sufficient generation of thrombin, the major enzyme of coagulation that plays a pivotal role in clot formation. The plasma concentration of FVIII defines clinical categories of hemophilia A as mild (> 5%), moderate (1-5%), or severe (< 1%) but within these categories, individuals with similar plasma levels can have different bleeding phenotypes (1). Some variations in bleeding phenotype can be assigned to the different mutations in the *F8* gene, or thrombophilic mutations, but a large portion remains unexplained(2, 3). Plasma protein levels are potential modifiers of bleeding; in particular, the variability in coagulation factor levels is quite large, with the normal range generall regarded as 50-150% of the mean of the healthy population. We hypothesize that certain combinations of coagulation factor levels within this normal range could enhance thrombin generation in the context of FVIII deficiencies and thus reduce bleeding. Identifying such combinations using a reductionist approach alone is unlikely to succeed since clot formation is a complex, nonlinear process. In this study, we use a mechanistic mathematical model of flow-mediated coagulation as a screening tool to predict modifiers of thrombin generation in FVIII deficiency and verify these predictions with experimental models.

The coagulation reaction network shares many features with gene, metabolic, and protein networks, for which mathematical and computational approaches are essential to decipher behavior and predict system responses (4–6). First, complex networks often display nonlinear responses due to the presence of positive and negative feedback loops. In the coagulation network, thrombin both enhances and inhibits its own production through different pathways. Second, the interactions between components of complex networks must be fully described to mechanistically explain emergent properties of the network itself. For example, our previous mathematical models of coagulation under flow showed that thrombin generation had a threshold dependence on the amount of exposed tissue factor (TF) (7–9), a prediction later validated in experiments (10). Finally, complex networks are robust in that they maintain phenotypic stability in the face of perturbations. Even with the normal variability in coagulation factors levels, the healthy hemostatic response is quite robust, leading to clots that prevent bleeding while maintaining vessel patency. The robustness of the coagulation network response to perturbations under disease states such as hemophilia is unknown.

A powerful tool for analyzing the variability of a model network’s output is sensitivity analysis (SA); here, model inputs are altered, either one-at-a-time (local SA, LSA) or in combination (global SA, GSA), and the resulting influence on model outputs is studied (11). In variance-based GSA methods, the variance in model output is decomposed and attributed to individual parameters and interactions between groups of parameters. One SA approach is to vary model parameters to identify those to which the model output is the most sensitive. There are LSA studies that use this approach on mathematical models of coagulation in the absence of flow (12, 13). Another approach is to use the model to make predictions about potential outcomes given variations in the input. For example, the inputs could represent variability in a disease state or to predict therapeutic outcomes or targets (14, 15). We previously conducted a LSA and GSA to determine how variation (±50%) in plasma levels of coagulation factors affected thrombin generation in a model of flow-mediated coagulation under healthy conditions (16). Our analysis revealed low overall variance of thrombin output, which is in line with results from Danforth et al. (15). Collectively, these results underscore the robustness of thrombin generation under healthy conditions.

In this study, we are interested in FVIII deficiencies and thus, we have fixed the FVIII level to be low in our mathematical model and performed a GSA by varying the remaining plasma protein levels. We used the GSA as an initial screening tool to search for combinations of plasma protein levels that either enhance or reduce thrombin generation in the context of FVIII deficiency. Combinations identified with the GSA to have the greatest effect were verified in whole blood flow assays and calibrated automated thrombography (CAT). This systems biology approach identified a potential mechanism where variations in FV levels within the normal range dramatically alter thrombin generation and fibrin formation in FVIII deficiencies.

### Brief overview of clotting

Blood clot formation involves the coupled processes of platelet aggregation and coagulation, which are triggered when blood is exposed to the subendothelium (SE). Platelet aggregation begins when platelets adhere to SE matrix proteins, become activated and form a platelet plug to arrest blood loss. Coagulation consists of a biochemical network that is initiated by TF, progresses by means of enzymatic reactions on activated platelet surfaces (APS) (17, 18), and culminates in thrombin generation. Thrombin activates platelets and converts the soluble fibrinogen into insoluble fibrin, which polymerizes to form a stabilizing mesh surrounding the platelet mass.

Coagulation proteins include inactive enzyme precursors (zymogens) factors VII, IX, X, XI and II (prothrombin) and the corresponding active enzymes factors VIIa, IXa, Xa, XIa, and IIa (thrombin), as well as the inactive/active cofactor pairs factors V/Va and VIII/VIIIa. Enzyme inhibitors include antithrombin (AT), tissue factor pathway inhibitor (TFPI), and activated protein C (APC). Thrombin generation occurs through the activity of three major cofactor/enzyme complexes, TF:FVIIa, FVIIIa:FIXa (tenase), and FVa:FXa (prothrombinase); each requires a suitable cellular surface on which to form, the SE for TF:FVIIa and APS for FVIIIa:FIXa and FVa:FXa. Coagulation proteins bind to specific binding sites on APS prior to forming platelet-bound complexes. We denote platelet-bound species with a prefix “plt”, e.g., plt-FV. Below we use “plasma proteins” to refer collectively to zymogens, inactive cofactors, and inhibitors.

The backbone of the coagulation reaction network involves the following steps (see Fig. S1A) that can greatly amplify the initiating signal of TF exposure: TF:FVIIa activates FIX and FX on the SE; FIXa and FXa bind to APS; plt-FXa activates small amounts of plt-FV and plt-FVIII; plt-FVIIIa and plt-FIXa form tenase complexes on APS; plt-FVa and plt-FXa form prothrombinase complexes on APS; prothrombinase converts prothrombin into thrombin. The backbone is augmented with numerous feedback loops which are critical to robust thrombin generation. For example, plt-Xa produced by tenase activates plt-VIII allowing more tenase to form; thrombin activates plt-FVIII and plt-FV allowing more tenase and prothrombinase to form. FVIII deficiencies reduce thrombin generation because less tenase forms on APS. Factor XII and blood-bourne sources of TF are not considered in our mathematical model.

## Results

The mathematical model we use is presented in (9, 16) and is an extension of our earlier models (7, 8, 19). The model includes all of the reactions depicted in Fig. SI1, with the equations and parameter choices fully described in (9). The inputs to this model are the TF level, plasma protein levels, platelet count, binding sites on APS, and flow rate; the available outputs are all model species’ concentrations as a function of time.

### Tissue factor density distributions for normal and FVI-II-deficient plasma

Our previous studies revealed a critical TF level, where thrombin sharply transitioned between an attenuated and amplified response (7–9). Under FVIII-deficient conditions and high TF, our model produces more than 1 nM of thrombin by 10 min, albeit at a decreased rate compared to normal FVIII levels (7). Furthermore, it is known that individuals with FVIII deficiencies bleed mostly in regions of the body with low TF levels (20–22). Taken together, these studies show that the TF level has an important influence on whether substantial thrombin is produced for any given values of plasma protein levels. Thus, to identify modifiers of hemophilia A, we first determine a range of critical TF levels so that minor decreases in that level result in little or no thrombin generation and minor increases result in substantial thrombin generation. As an initial screen, we varied the plasma protein levels between 50 and 150% of their physiologic levels for both normal and FVIII-deficient plasma (FVIII fixed to 1% or normal), using 2,500 Latin Hypercube samples (23). For each of these parameter set samples, a bisection procedure was used to determine the minimum TF level required to achieve an amplified thrombin response, i.e., 1 nM thrombin by 40 min. The distribution of these values determines a critical range of TF levels over which the thrombin response is most sensitive to variations in plasma protein levels. In our model, thrombin that reaches 1 nM (indicated by a blue line in Fig. 1B) activates platelets and is then likely to continue to increase (7–9, 16).

**Fig. 1.**
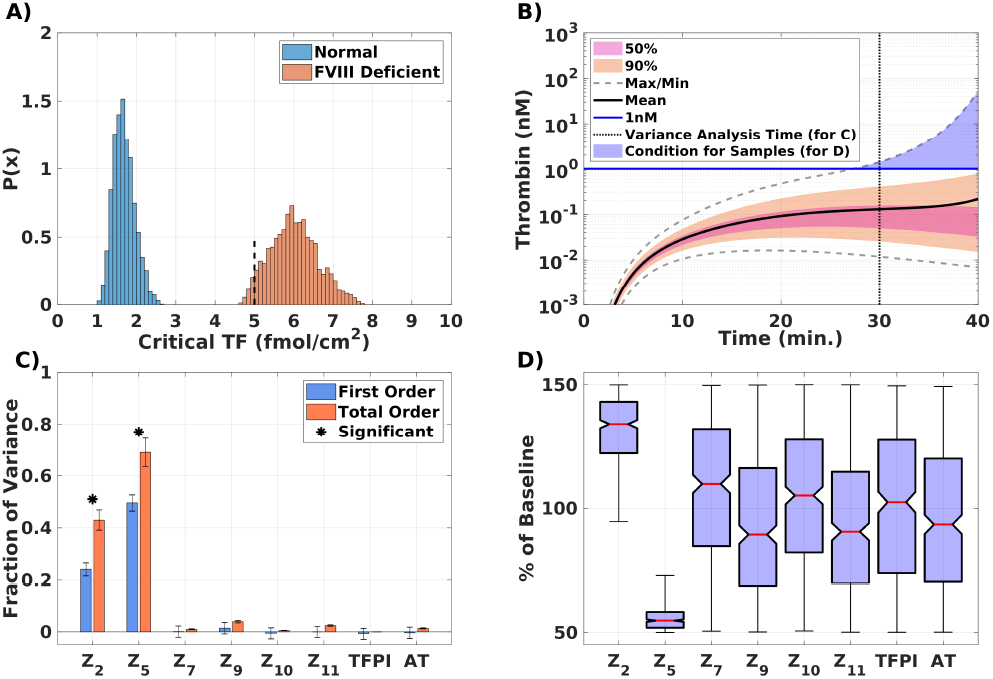
Global sensitivity analysis for mathematical model of flow-mediated coagulation. **A)** Critical TF distributions for normal and FVIII deficient plasma. Dashed black line at 5 fmol/cm^2^ is the fixed TF for all further simulations. **B)** Thrombin concentration time series generated by uniformly and independently varying plasma protein levels ±50% from normal (110,000 total simulations); mean (solid black line), boundaries that encompass 50% (pink), and 90% of the data (orange), and the maximum/minimum of the computed solutions (gray-dashed); blue line drawn at 1nM. **C)** First (blue) and total (orange) order Sobol indices are plotted as mean ± standard deviation computed with 5,000 bootstrap samples of the original 110,000 simulations. **D)** Plasma zymogen and inhibitor levels distributions shown as box-and-whisker plots (mean in red, whiskers *3IQR*), conditioned on achieving more than 1nM of total thrombin.

Fig. 1A shows the TF distributions for the normal (blue) and FVIII-deficient (orange) plasma where the protein levels were varied between 50-150%. No overlap between the two distributions is observed, with normal and FVIII-deficient plasma having a TF range of [1.07, 2.62] fmol/cm^2^ and [4.63, 7.78] fmol/cm^2^, respectively. These distributions suggest that a TF level of 5 fmol/cm^2^, near the left edge of the distribution for FVIII-deficient plasma, is a good choice for conducting further probes of plasma protein levels in a GSA, because for the majority, but not all, of the plasma protein combinations that we tested, little thrombin is produced. All further simulations in this study are performed with TF at 5 fmol/cm^2^.

### GSA identifies FV as a modifier of thrombin generation in hemophilia A

We performed a GSA of thrombin generation by varying plasma protein levels using 110,000 samples in which each plasma protein level (except FVIII, which was fixed to 1%) was sampled uniformly and independently between 50-150% of normal. Quantiles of the thrombin concentration time-course are shown in Fig. 1B. While no simulation achieved more than 100 nM by 40 min, about 5% of the simulations eventually led to thrombin greater than 100 nM (not shown). We further distinguish simulations by those that led to 1 nM thrombin within 40 min and those that did not.

To assign fractions of the thrombin output variance observed in the quantiles of Fig. 1B, we performed a Sobol analysis (24). Fig. 1C shows the first order (blue) and total order (orange) Sobol sensitivity indices for the thrombin concentration at 30 min. The first order indices represent the fraction of the variance attributable to that one parameter alone while the total order index indicates the fraction due to that parameter plus its interactions with other parameters. We see that the parameters, i.e., the plasma protein levels, that had the most influence on the variance in thrombin at 30 min are FV and prothrombin (FII), which account for approximately 50% and 24% of the variance, respectively. In addition, approximately 19% of the model variance is explained by the interaction between prothrombin and FV, as indicated by the total order Sobol indices exceeding the first order indices.

Next, we characterized the ≈ 5000 simulations that led to 1 nM thrombin within 40 min (blue shaded region in Fig. 1B). Fig. 1D shows the distribution of all plasma proteins that correspond to those simulations. The majority of the plasma protein levels have medians close to their average and distributions that appear roughly uniform. This indicates that 1 nM thrombin is possible with any value of these plasma levels. The plasma FV and prothrombin levels were striking exceptions. Both have medians close to their extreme values and are distributed over a narrower range than the other plasma protein levels. Although the prothrombin distribution is skewed towards higher levels, it does extend below 100%, indicating that higher than normal prothrombin is not strictly necessary to achieve 1 nM thrombin. Conversely, every sample that achieved 1nM thrombin had plasma levels of FV that were strictly less than 75% of normal. Indeed, over 75% of the samples had a FV level less than 60% of normal. This indicates that FV levels on the low end of normal (close to 50%) are necessary to enhance thrombin generation in FVIII-deficient plasma stimulated with a low level of TF in this model.

### Exploration of Mechanism: The competition for FXa

We used our mathematical model to explore how lower FV can enhance thrombin production in FVIII-deficient plasma. Table S1 lists the variations in plasma levels of prothrombin and FV that we considered. Fig. 2A shows the time-course of thrombin for these four cases. In the two cases with low FV, substantial thrombin is produced and thrombin generation occurs earlier with a higher prothrombin level. Very little thrombin is produced with normal levels of FV. These results indicate that low FV is key to enhancing thrombin production and increasing prothrombin level alone does not. Fig. 2B-C show that the tenase concentration associated with low FV diverges from normal FV as early as 5 min, before the prothrombinase and thrombin curves diverge. Given that there is little thrombin at these early times, the differences in the tenase behavior suggest that early competition between plt-FVIII and plt-FV to bind to plt-FXa plays a significant role. To see this competition, Fig. 2D-E shows the concentrations of plt-FVIII:FXa and plt-FV:FXa. The concentration of plt-FVIII:FXa is substantially higher for the low FV cases than for the normal FV cases, and for each FV level, there is very little difference (for at least 20 min) between the high and low prothrombin levels. To further demonstrate the importance of FV and FVIII competition for FXa, we adjust the kinetic rate constants for these reactions while setting the FV and FII concentrations to their baseline levels (see Fig. S2A-B). We found that: increasing only the rate of binding of plt-FVIII to plt-FXa leads to very low thrombin production; decreasing the rate of binding of plt-FV to plt-FXa by 50% and increasing the rate of binding of plt-FVIII to plt-FXa by 50% from their normal values, however, results in a 30-fold higher thrombin concentration at 40 min and thrombin generation reaches 1 nM at about 37 min. These results confirm that competition between plt-FV and plt-FVIII for plt-FXa influences the rescue of thrombin under FVIII-deficient conditions.

**Fig. 2.**
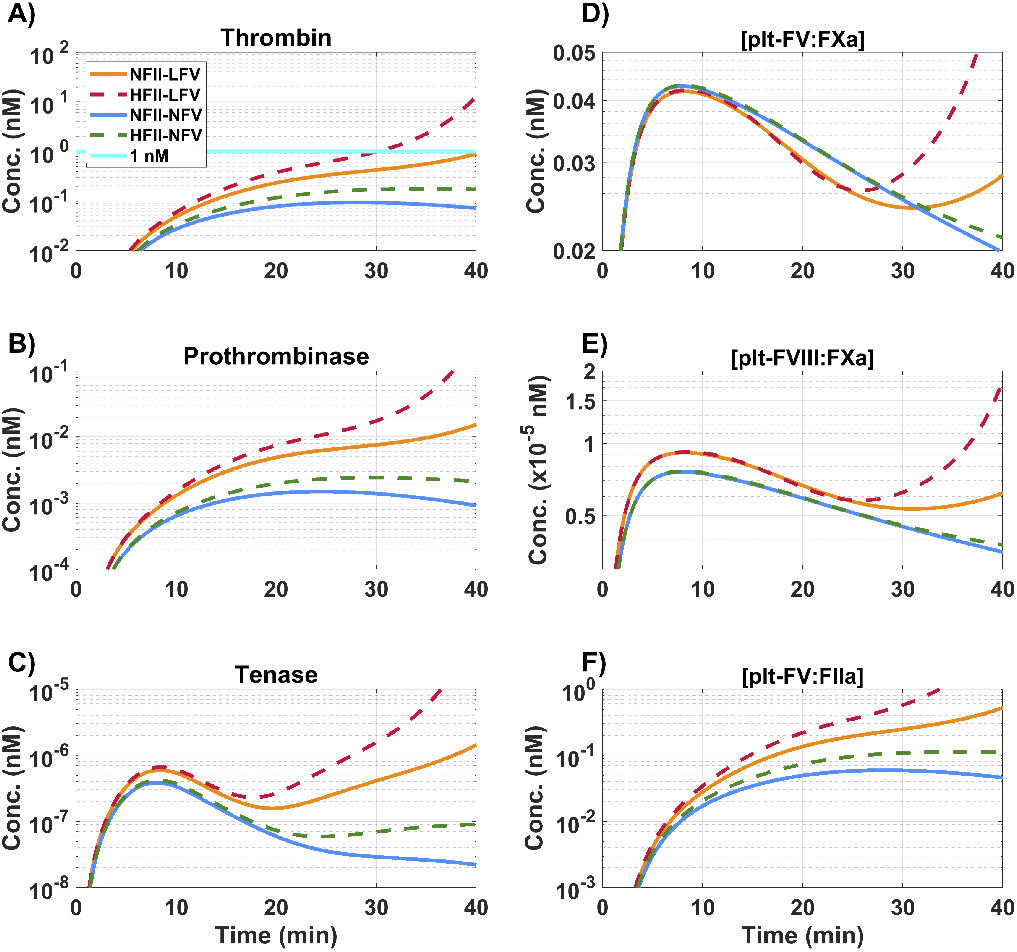
Effects of variations in plasma FII and FV levels on thrombin generation and the evolution of enzyme complexes in the mathematical model. “N” denotes 100%,”L” denotes 50%, and “H” denotes 150% of the respective baseline plasma level; “NFII” = 1400 nM, “NFV” = 10 nM. **A)** Total thrombin; **B)** prothrombinase (FVa:FXa); **C)** platelet tenase (FVIIIa:FIXa); **D)** FVIII:FXa complex on plt; **E)** FV:FXa complex on plt; **F)** FV:FIIa complex on plt; during a time course of 40 min. Description of labels found in Table S1.

Thrombin activation of FV and FVIII, and APC binding to FVa and FVIIIa, may also influence thrombin production in the low FV cases. To explore this, we performed simulations in which the binding rates for FV and FVIII to thrombin and for FVa and FVIIIa to APC were set to zero (Fig. S3A-F), and compared the results to those in Fig. 2A-F. We see that thrombin generation is initially increased in the low FV cases but is not sustained after 40 min (Fig. S3A). Tenase and prothrombinase follow a similar trend, where low FV produces higher concentrations initially but amplification of tenase and prothrombinase complex formation does not occur (Fig. S3B-C). Thus, thrombin-mediated activation of FV and FVIII is necessary to produce a significant thrombin response despite the lack of APC-mediated inhibition.

It is not intuitive how lowering FV plasma levels results in near-normal prothrombinase (see Fig. 2B) since FV is a precursor of a component of prothrombinase. In the low FV cases, despite decreased FXa-mediated activation of FV (Fig. 2D), the concentration of plt-FV bound to thrombin at 20 min is approximately 10-fold larger than that associated with normal FV cases (Fig. 2F). This is a direct result of the increased thrombin concentration shown in Fig. 2A. Thrombin feeds back by activating plt-FVIII and indirectly, plt-FIX via FXIa, to form more tenase (Fig. S4A-B). Increased tenase results in more plt-FXa, which binds to the thrombin-activated plt-FVa, leading to more prothrombinase and thus more thrombin, even with FVIII deficiency.

In summary, we used our mathematical model to identify where in the coagulation reaction network FV and FVIII most strongly interact. The coagulation network involves no direct reaction between FV and FVIII, but they compete for binding to FXa and thrombin. We confirmed that thrombin-mediated activation of FV and FVIII is essential to a substantial thrombin response. More importantly, the early divergence of thrombin and key complexes in low versus normal FV cases, even when thrombin-mediated activation of FV and FVIII is turned off, identifies the competition of FV and FVIII for FXa as the initiator of thrombin generation rescue in low FV, FVIII-deficient blood. Additional support for this hypothesis comes from simulations in which we isolated the reactions amongst plt-FV, plt-FVIII, and plt-FXa by varying their binding rates.

### Partial inhibition of FV enhances fibrin deposition in FVIII-deficient blood under flow

Whole blood microflu-idic assays were performed at 100 s^−1^ on type I collagen-TF (1.09 ± 0.2 fmol/cm^2^). Blood from individuals with moderate and mild FVIII deficiencies was treated with exogenous prothrombin (50 *μ*g/mL), an anti-FV antibody at a concentration that reduced FV activity to ~ 60% in normal pooled plasma (Fig. S5), both exogenous prothrombin and anti-FV, or a vehicle control (Figs. 3, S6). In the vehicle and exogenous prothrombin cases, little to no fibrin was observed. Platelet adhered and nearly coated the surface over the 25 min experiment, but there were were no large, multilayer platelet aggregates. Anti-FV alone supports fibrin formation in and around multilayer platelet aggregates, and when combined with prothrombin, the effect is even more pronounced with larger, platelet-fibrin thrombi forming. There was a significant increase in both the rate of and maximum accumulation of fibrin(ogen) with partial inhibition of FV. There were no significant differences in total platelet accumulation despite the morphological differences described above (Fig. S8). The FVIII levels in these samples were higher (3.0-8.5%) the day of the experiments than those considered in our mathematical model (1% FVIII) because these individuals have mild to moderate FVIII deficiencies (SI Appendix). Nevertheless, the inhibition of FV clearly shows an increase in fibrin deposition in these experiments, and indirectly, thrombin generation.

**Fig. 3.**
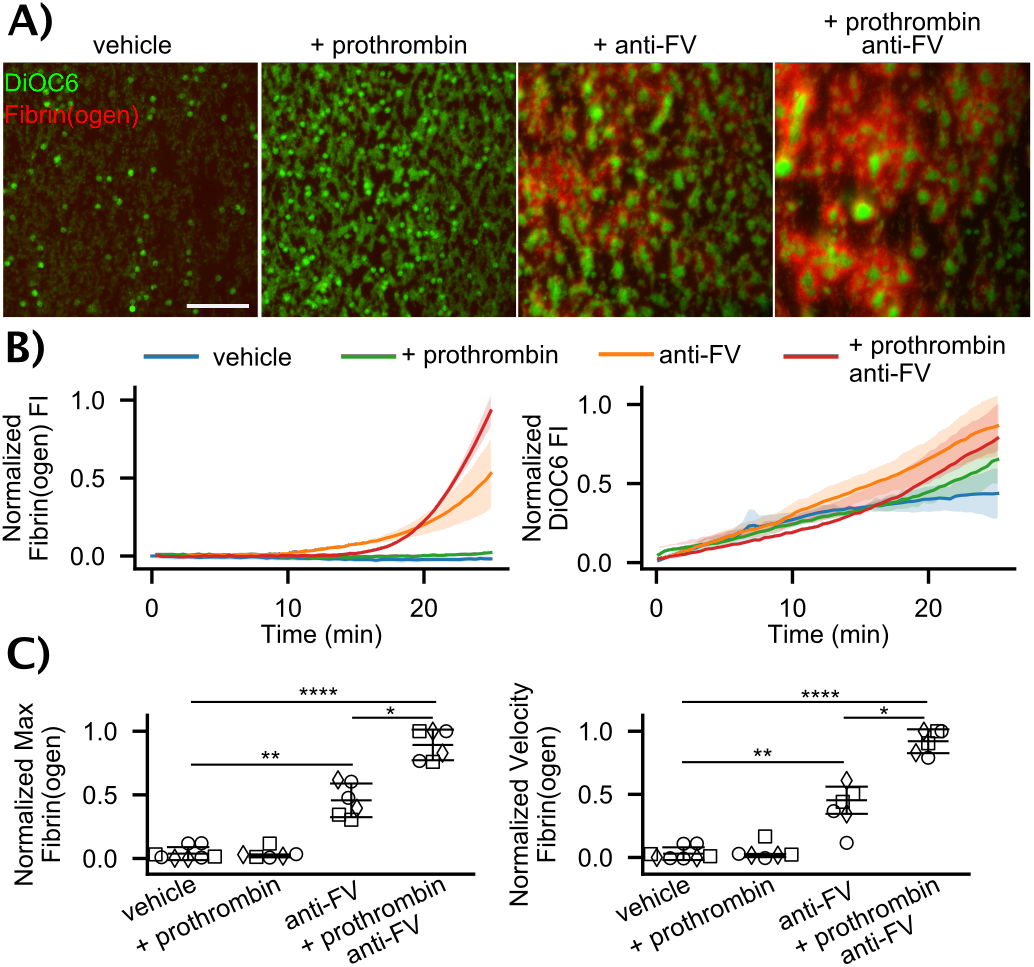
Flow assays with whole blood from FVIII deficient individuals. **A)** Representative images of DiOC6 labeled platelets and leukocytes and Alexa Fluor 555 labeled fibrin(ogen) on collagen-TF surfaces at 100 *s*^−1^ after 25 min for vehicle control, 50 *μ*g/mL exogeneous prothrombin, 100 *μ*g/mL anti-FV, and exogenous prothrombin and anti-FV. Scale bar = 50 *μ*m. Individual fluorescent channels are found in Fig. S6. **B)** Representative fibrin(ogen) and platelet/leukocyte accumulation dynamics in terms of normalized fluorescent intensity (FI). **C)** Fibrin(ogen) normalized maximum fluorescence intensity and rate of deposition (normalized velocity) for FVIII levels of ◯ = 3.0%, ◸ = 7.5%, ◊ = 8.5%. See SI and Fig. S7 for calculation of metrics. P-values represented as *, **, and **** for 10^−2^,10^−4^, and 10^−7^, respectively.

### Low FV and partial inhibition of FV enhances thrombin generation in FVIII-inhibited or FVIII-deficient plasma

We used calibrated automated thrombography (CAT) (25) to measure the effects of reducing FV levels or activity on thrombin generation dynamics in a clinical clotting assay. We altered zymogen concentrations to approximate the four conditions in Table S1 using mixtures of FV and prothrombin depleted plasmas, purified FV and prothrombin, and an anti-FVIII function blocking antibody used at a concentration that yields FVIII activity of <1% to simulate severe hemophilia A. Consistent with mathematical model predictions, Fig. 4A and Table S3 show that low FV (43%) increases the peak thrombin concentration, which is further enhanced when prothrombin (136%) is added. Similar trends, albeit with lower peak thrombin concentrations, were measured with FVIII deficient plasma using the same treatments as the flow assay experiments described in the previous section (Table S4). Fig. 4B shows results from an individual with severe FVIII-deficiency (< 1%) and partial inhibition of FV (65% activity) increases thrombin peak concentration, with an even larger peak concentration when combined with high prothrombin (135%). Notably, there is an increased lag time for treatments including the anti-FV antibody in FVIII-deficient plasma. This observation could be due to the difference between inhibiting FVIII with an antibody compared to plasma deficient in FVIII.

**Fig. 4.**
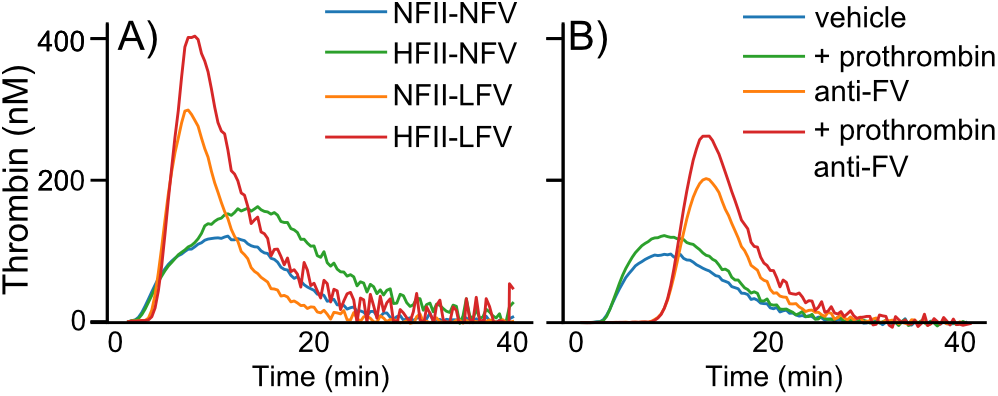
Calibrated automated thrombography. A) FII and FV levels were varied using immunodepleted plasmas and purified FII and FV in the presence of an anti-FVIII function blocking antibody. ‘N’, ‘H’, and ‘L’ corresponds to normal, high, and low levels of respective zymogen. B) FVIII deficient (< 1%) plasma treated with vehicle control, 50 *μ*g/mL exogenous prothrombin, 100 *μ*g/mL anti-FV, and exogenous prothrombin and anti-FV. All assays conducted with 5 pM TF and phospholipids. Tables S3 and S4 contain the measured prothrombin, FV, and FVIII levels corresponding to each curve.

## Discussion

In this study, we showed that a mechanistic mathematical model of flow-mediated coagulation can identify important modifiers of network dynamics. We were motivated by observations of the variability in bleeding among individuals with hemophilia A with similar FVIII levels. We hypothesized that variations in coagulation plasma protein levels within the normal range could modify thrombin generation when FVIII is outside the normal range. We determined critical TF levels necessary for the model to produce 1 nM thrombin by 40 min; this was meant to represent a TF level where bleeding would be common in hemophilia A but where significant thrombin generation is still possible. Using GSA on the model at the specified TF level, we identified that FV levels at the low end of the normal range could push the coagulation system to generate a significant thrombin response. When this level of FV was combined with prothrombin levels at the high end of the normal range, thrombin generation was enhanced even further. This prediction was verified in a microfluidic model of thrombus formation on collagen-TF where fibrin accumulation was used as a proxy for thrombin generation, and in thrombin generation assays using immunodepleted plasma to match the plasma composition of the mathematical model and in plasma with a severe FVIII deficiency. Exploration with the model revealed a potential mechanism to explain these observations; a reduction in FV frees plt-FXa to activate plt-FVIII, leading to more tenase and more prothrombinase on APS, which ultimately boosts thrombin generation. The modeling results in this paper depend on the assumption that plt-FXa can activate plt-FV and plt-FVIII. In our simulations, plt-FXa is the dominant activator of FV and FVIII before significant amounts of thrombin have been produced. The assumption that FXa can activate FV on an activated platelet’s surface is based on data of Monkovic and Tracy (26) and studies using tick saliva protein (27). In regards to activation of FVIII by FXa, there is in vitro evidence that FVIII bound to an APS can be activated by both thrombin and FXa (28, 29). Because FVIII circulates in the plasma bound with von Willebrand factor (VWF) and while bound may be protected from activation by FXa (30, 31), some have suggested that FXa-mediated activation of FVIII occurs to a minimal extent in vivo. This protection may be mitigated because during clot formation vWF binds to APS and the FVIII attached to this vWF may redistribute to the APS, given that FVIII has similar affinities for VWF and phopholipid surfaces (32, 33). Our results suggest that further biochemical studies of FXa’s role in activating FVIII are needed.

An alternative view of the early stages of coagulation is that a trace amount of thrombin, produced on TF-bearing cells, is responsible for initial FV and FVIII activation (17, 34). This view comes, in part, from experiments in a static cell-based model of coagulation (17) in which initial platelet activation itself seems to rely on this thrombin as there is no collagen exposure in that system. If translated to a situation of vascular injury under continued flow (a situation that our mathematical model and flow assay simulate), the trace thrombin would be produced on the vascular wall and could activate FVIII and FV in the plasma or on APS. Under flow conditions, even at the low shear rate of 100 s^−1^ used in our model simulations, this view is problematic. Our model predicts that ≈ 99% of the FXa and FIXa produced by TF:FVIIa on the vascular wall is quickly washed away by the flow and does not reach the surfaces of activated platelets (8). The same would be true of any thrombin produced on the vascular wall and so, no more than 1% of any trace amounts of this thrombin would make it to activated platelets in order to carry out its putative activation of FV and FVIII there.

FV is contained in and secreted from platelet *α*-granules, a mechanism that is incorporated into our model (7, 9). Approximately 20% of the FV in blood is contained in the platelets (35). Platelet FV comes from plasma FV that is endocytosed by megakaryocytes (36–38), thus we assume in the model that a percent change in FV plasma levels correlates with an equal percent change in the platelet FV levels. However, it is unknown what the true correlation is between the two FV pools. Platelet FV is distinct from plasma FV due to modifications in megakaryocytes (39) and is more procoagulant than plasma FV (36, 40). In our model both FV pools have the same biochemical characteristics. Future work is needed to tease out the relative roles of plasma and platelet FV in thrombin generation in the context of hemophilia A.

We are unaware of previous reports demonstrating a relationship between FVIII deficiencies, FV levels within the normal range, and thrombin generation. A mutation in a molecular chaperone that transports proteins from the endoplasmic reticulum to the Golgi results in a combined FV and FVIII deficiency (41). This mutation causes low levels (5-30%) of both FV and FVIII. That situation is different than our findings where thrombin generation is modulated by FV levels within the normal range (50-150%) for FVIII deficiencies. There are reports of individuals with both a FVIII deficiency and a common variant of FV called FV Leiden (rs6025) (42). FVa’s endogenous inhibitor, APC, cannot bind to this variant, leading to a hypercoagulable state. Individuals with combined FVIII deficiency and FV Leiden have a milder bleeding phenotype (43), but this is distinct from the effect we show here where reduced FV levels allow for more FVIII binding to FXa on APS.

There are several potential implications for our findings. FV levels could be an inherent modifier of bleeding risk in combination with FVIII deficiency in hemophilia A. Studies of clinical bleeding in individuals with hemophilia A are needed to support this hypothesis. Our results also suggest that temporal changes in FV expression, such as those related to circadian rhythms or menstrual cycle, could influence bleeding risk. Finally, our study shows that a systems biology approach to coagulation may facilitate the discovery of previously unrecognized interactions between the several components of the system and may serve as a platform to study other highly complex clinical problems.

## Methods

All human subjects protocols were approved by the Institutional Review Board of the University of Colorado, Denver I Anschutz Medical Campus. Details regarding the mathematical model description, implementation, and parameter choices, and materials and methods for flow assays and CAT are found in the SI Appendix, SI Materials and Methods.

## ACKNOWLEDGEMENTS

This work was supported in part by National Institutes of Health (R01 HL120728) and National Science Foundation (CBET-1351672)

AUTHOR CONTRIBUTIONS
KBN, KL, ALF, SSS, JDP designed research. KGL, MTS, MGS, MB, KR, MJMJ performed research. KBN, KL, ALF, SSS, JDP, KGL, MTS, MGS, MB wrote the paper.

